# A strong relationship between environmental DNA metabarcoding and rank-based abundance of fish

**DOI:** 10.1101/2025.01.22.634322

**Authors:** Joanne E Littlefair, Lauren D Hayhurst, Matthew C Yates, Michael D Rennie, Melania E Cristescu

## Abstract

1. Increasingly, molecular methods of species monitoring are integrated into freshwater biodiversity surveys and fisheries management. Inferring organism abundance or biomass from sequence counts derived from metabarcoding data has been an exciting but contentious concept in the biomonitoring community for some time. Although demonstrating a strong correlation with abundance has proven difficult, many researchers have assumed that quantitative metabarcoding data can at least provide broad-scale ranking of abundance. However, robust field validations of this widely-held assumption remain scarce.
2. Here, we analyse metabarcoding read counts of fish eDNA data derived from 20 lakes and use betabinomial mixed effects models to compare this to rank abundance generated from long-term fish survey data. Rank abundance data for 18 species was generated within-species across-sites, meaning that ranks compare the abundance of the same species in different lakes. We also investigated a possible allometric effect on eDNA production by analysing a subset of data for effects of fish body mass on the amount of eDNA sequences.
3. We found a good relationship between species-specific eDNA sequences and within-species rank abundance categories for fishes, with rare fish producing 3% of sequences in a library, moderately abundant producing 7% and abundant fish producing 29%, according to model predictions.
4. We found a small negative effect of body mass on the amount of eDNA sequences, where the proportion of reads recovered significantly decreased with increased mean body mass of the population.
5. *Synthesis and applications:* The benefit of this approach is the potential for rapid assessment of rank abundance for multiple species, including smaller species which are often missed by conventional methods such as gillnetting, with relatively low amounts of additional effort. This approach will assist practitioners taking a species-based approach to freshwater habitat management in lakes worldwide.

## Introduction

Abundance data are critical to understanding many of the fundamental concepts in ecology and evolution, including population dynamics, community ecology, and macroecology (McGill & Collins, 2003; Preston, 1948). At the local scale, abundance underlies some of the few “laws” of community ecology, such as species-abundance distributions (McGill et al., 2007). From an applied perspective, abundance is a key component of biomonitoring schemes used to understand the functionality of ecosystems and plan effectively for ecosystem management. Abundance data is integrated into many biomonitoring schemes mandated by national and international legislation; including the EU Habitats Directive and the US Endangered Species Act (Evansen et al., 2021; Moussy et al., 2022). Abundance data can take multiple forms but aims to capture information about the number of individuals in a species within a particular area (Begon et al., 2006). Ideally, populations are measured in absolute numbers (often estimated directly through observational methods such as mark- recapture, spatial capture-recapture, or distance sampling methods), but these methods are often intensive, invasive for the species studied, and require significant effort. More commonly, abundance is expressed relative to some standardized capture method (often termed ‘relative abundance’ and expressed as catch per unit effort, CPUE). This can be based on factors such as camera-trap hours, amount of gear used, deployment duration, or transect length.

Environmental DNA (eDNA) techniques are increasingly being used alongside conventional monitoring techniques by academics, industry, and NGOs for species monitoring due to their rapid, large-scale and non-invasive nature (Deiner et al., 2017). Initially, eDNA studies focussed on reporting species presence/absence status, but multiple reviews have emphasised the potential for integrating abundance data into eDNA research (Deiner et al., 2017; Yao et al., 2022; Zaiko et al., 2018). Furthermore, this is a common enquiry from end-users interested in using eDNA as a species monitoring technique. Two kinds of abundance metric have been investigated using environmental DNA data (Figure 1; Luo et al., 2022). Within- species across-site abundance compares the relative abundance of the same species at multiple sites (e.g., qPCR analysis of eDNA from a single species at different sites within a single river or different rivers). The second abundance metric concerns within-site across- species measurement in which the relative abundance of different species in a single habitat (e.g., one lake or river) are compared. This is most often used in metabarcoding studies, which compare eDNA sequences to relative rank abundance of species within one site (Hänfling et al., 2016; Lawson Handley et al., 2019; Sard et al., 2019).

**Figure 1:**
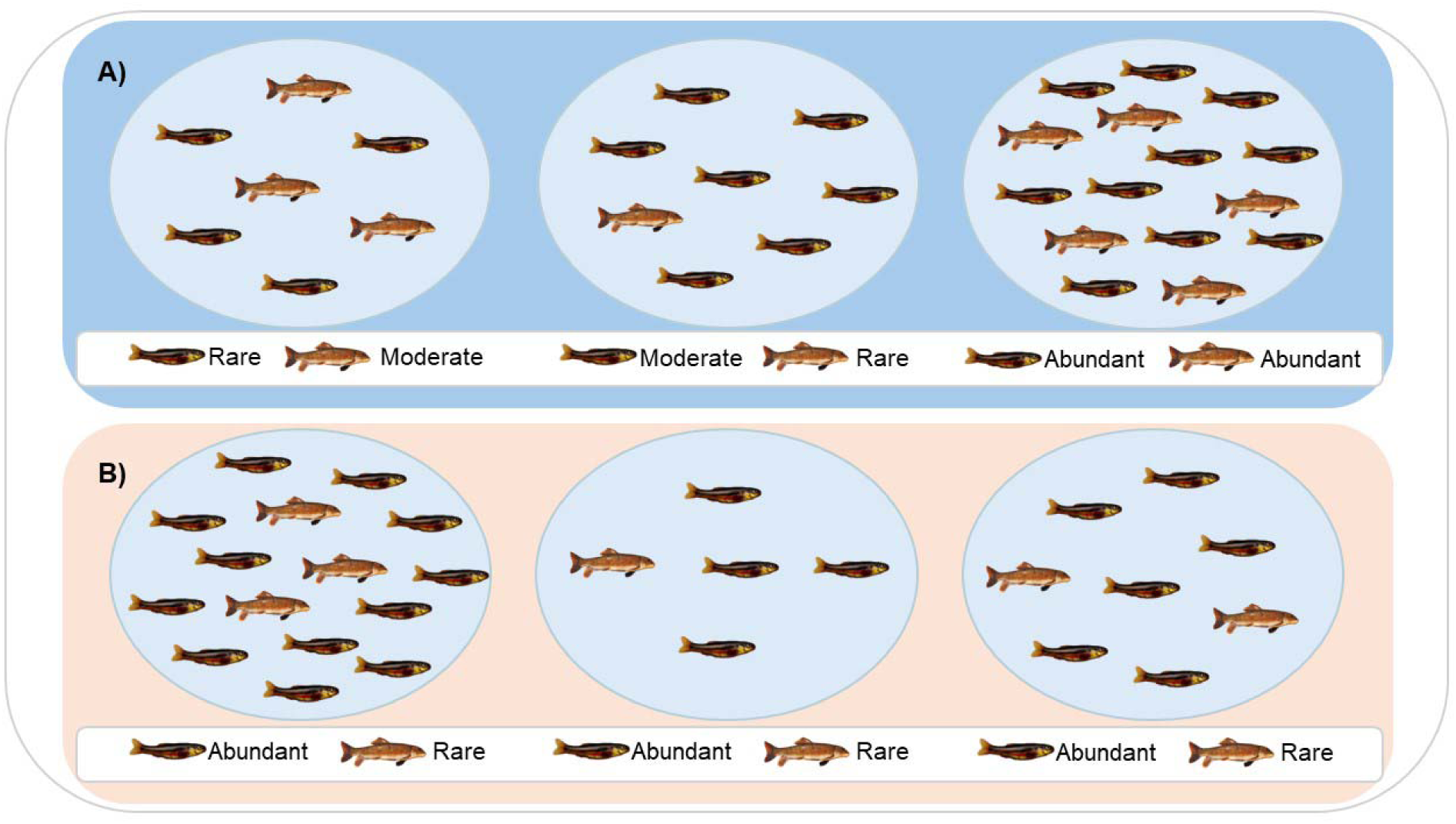
Conceptual diagram of across-lake, within-species rank abundance (A), compared with within-lake, across-species rank abundance (B).

In fisheries, abundance data provide critical information for resource practitioners to monitor and manage exploited stocks, e.g., through setting quota allotment for commercial fisheries or harvest regulations around sport fisheries. Management of sport fisheries (e.g., of lakes on a landscape) often relies on the use of CPUE from standardized collection methods (e.g., North American Standard, NORDIC, etc; Morgan & Snucins, 2005; Sandstrom et al., 2018), where the abundance of fish is comparable to the catch from other lakes using the same collection methods. Within-species across-site abundance (for multiple species across multiple habitats) is directly comparable to CPUE obtained by traditional standardized collection surveys used by fisheries managers to determine community composition and estimate the relative abundance of fishes across comparable waterbodies on the landscape. While many agencies are developing calibration equations to convert relative abundance to absolute abundance estimates (e.g., by applying these standardized capture methods to populations that have existing mark-recapture information), in the absence of such information, CPUE abundance data provides essentially rank-based information on the number of fish of a particular species in one system versus another, relative to all other systems sampled by the same sampling methods.

Within-site across-species abundance is used within single habitats especially in the context of food web ecology to understand dominant energy pathways within the ecosystem (Jepsen & Winemiller, 2002) and cascading effects of environmental disturbance (Kidd et al., 2014). Fish are important predators within lake ecosystems and a change in the abundance of a particular species can result in cascading effects on the whole community. This can be in the context of management of a single lake; for example, culling of a fish species to manage overcrowding or parasitic infection (Amundsen et al., 2019), or studying the consequences of species invasion (Gallardo et al., 2016). Changes in ecosystem structure and the effects of interactions between target and non-target species form the basis of ecosystem-based fisheries management, which aims to move beyond single-species assessment to understanding the effects of management decisions for a particular target fish species on the entire ecosystem (Mangel & Levin, 2005).

To date, researchers have primarily investigated the relationship between eDNA concentration (e.g., copy numbers/L) generated by qPCR or ddPCR data, and a measurement of density or biomass generated from mark-recapture or relative measures of abundance or biomass from standardised trap or net data. Often, studies and meta-analysis have found positive relationships between these variables (Lacoursière-Roussel et al. 2016; Spear et al., 2021; Thalinger et al., 2019; Tillotson et al., 2018; Yates et al., 2019), although factors such as strong currents in dynamic freshwater environments (Barnes & Turner, 2016; Rice et al., 2018; Tillotson et al., 2018), animal behaviour and reproductive status (Doi et al., 2016) can contribute to a weaker correlation. Metabarcoding derived data has a more tenuous relationship with organism abundance or biomass. A meta-analysis of mock communities created with either biomass, nucleic acids or individual whole-organisms as the input material showed a weak relationship with the percentage of output sequences (Lamb et al., 2019). This study reported a high degree of uncertainty associated with this result due to a large amount of inter-study variation. Scaling up from mock communities, the ultimate goal is to use environmental DNA sequences to inform population abundance estimates in natural habitats. Within single waterbodies, metabarcoding read count or site occupancy has been correlated with across-species rank abundance derived from long-term datasets, expert opinion and catch composition in marine fisheries (Green et al., 2024; Hänfling et al., 2016; Lawson Handley et al., 2019; Sard et al., 2019). Despite these correlations within-site across-species, validation of within-species across-site abundance using metabarcoding eDNA remains elusive.

To address this knowledge gap, we tested the relationship between sequence counts derived from metabarcoding data and estimates of **within-species across-site** rank-based abundance derived from conventional fishing techniques across multiple species in 20 lakes.

## Methods

Existing eDNA datasets were used in this analysis which were derived from sampling conducted in June-July 2017 at the International Institute for Sustainable Development Experimental Lakes Area (IISD ELA), northwestern Ontario, Canada (Littlefair et al. 2023b, Supporting Information). Water samples were collected from 20 lakes ranging in size from 2- 210 hectares and analysed for fish eDNA with metabarcoding (methods are summarized in the Supporting Information). One lake (lake 653) was removed from the initial dataset as it did not have associated rank-abundance data for fish.

### Assessment of rank-based fish abundance

A general metric of relative abundance (besides mark-recapture and catch-per-unit-effort estimates) has been implemented at the IISD ELA since its inception, originally proposed by Beamish et al., (1976). This assigns qualitative descriptors of 0 (absent), R (rare), M (moderately abundant) and A (abundant) to fish species across lakes. This was initiated during the first surveys for fishes performed in 1972-73 at the IISD ELA using gillnets, trapnetting and minnow trapping, described in Beamish et al., (1976). Since then, populations have been assessed (i.e. one to five times) using this methodology to provide information on fish communities, or on an annual or semi-annual basis in intensively sampled lakes (e.g. LTER and experimental lakes) using non-destructive methods. Crucially, the qualitative descriptors used in this method are applied within a species across lakes (i.e. Figure 1A) so that the absolute number of fish in each category varies for different species.

The precise means by which Beamish set these categories was not clearly documented, and therefore required validation using quantitative measures of abundance collected from these lakes since these original surveys. For this study, we performed additional validation comparing the qualitative descriptors initiated by Beamish to quantitative measures of abundance collected from routine IISD ELA sampling (i.e. catch-per-unit-effort or mark-recapture estimates; see Supporting Information).

### Statistical analysis

We examined the relationship between eDNA and species abundance by comparing species-specific read counts with fisheries-based rank abundance. Species-specific read counts, expressed as a proportion of total per-sample library size, was included as a response variable and modelled using a zero-inflated beta-binomial distribution with the response variable weighted by total library size. Beta-binomial distributions have been proposed as a candidate strategy for modelling microbial relative abundances in overdispersed data with a high proportion of zero values (Martin et al., 2020). eDNA samples returning an overall read count of zero were removed from the analysis as they caused convergence failure in early tests.

Random intercept effects for Lake, Sample ID and fish species were included in the statistical model to account for non-independence among samples and sites. The species random effect was also conditioned on rank-based abundance to reveal across-species patterns in read counts with rank abundance. Fisheries-based rank abundance data were fitted as a categorical fixed effect. An overdispersion term conditioned on rank abundance was included in the model, and rank abundance was also included as an explanatory variable in the zero-inflated component of the model. Simulated model residuals were examined using DHARMa v0.4.4 (Hartig, 2021). Model terms were evaluated using a nested backwards stepwise procedure based on AIC (Akaike, 1974). Overdispersion terms were tested first, followed by zero-inflated terms, random effects (Species conditioned on rank-based abundance), and fixed effects. However, all intercept-level random-effects were retained regardless of model AIC due to the non-independent sample design. All analyses were performed using the R package glmmTMB v 1.1.2.3 (Brooks et al., 2017) in R v 4.3.1 (R core team 2023).

We also evaluated the impact of body-mass on eDNA sequences. While large fish generally produce more eDNA than smaller fish, large fish may produce less eDNA on a per-gram basis (Maruyama et al. 2014, Yates et al. 2021). To investigate this, we (re)analysed a subset that included species for which mean body mass estimates were available. In addition to the full model above, mean body mass for each population in a lake was included as a continuous fixed effect. Mean body mass was either based on observed data or estimated from length-weight relationships (Table S1). If a species lacked body mass data for a subset of the lake systems in the dataset, mean body mass across lakes included in the current study was used. For a single species (Troutperch), mass data were unavailable; we therefore used mean mass data from other lakes at IISD ELA as a proxy for body mass in our study lakes. All species marked as ‘absent’ in traditional fisheries surveys were excluded from these analyses, given that no body mass data were available. Model terms were tested using stepwise backwards model selection based on model AIC.

## Results

The sequencing run returned 85,504,066 raw reads (mean 201,186 per sample) and produced 585 ASVs. Of these, 106 ASVs were assigned taxonomy at species level, and 87.1% of 12S sequences mapped onto ASVs corresponding to fish from the Lake of the Woods region in Ontario. Our eDNA analysis detected 23 fish taxa, nearly all identified to species level, except for two pairs of congeners: *Chrosomus eos* and *Chrosomus neogaeus* and *Coregonus artedi* and *Coregonus clupeaformis.* Rank abundance estimates for 18 of these species or genera were derived from conventional methods. Of the remaining five species, rank abundance estimates were not available because four of the species had rare distributions and one (*Esox masquinongy*) was a new detection in the largest lake in our analysis. Across the fisheries-based data, there were 115 assessments of rank abundance (rare=36, moderate=43, abundant=36), and 322 instances in which fish were assessed to be absent. After removing libraries with zero total sequences and lakes where there was no conventional fisheries-based assessment of abundance, we retained eDNA data from 386 samples for further analysis.

### Statistical analysis

All terms included in the model substantially improved the model’s fit, as indicated by ΔAIC >2, and were retained (Table 1). We calculated back-transformed model predictions to estimate the influence of the rank-based levels of abundance on the proportion of species-specific sequences within a library. For example, if species A was at rare abundance in a lake, our model estimated that 3% of sequences in a sample will belong to that species; if species B was at moderate abundance, our model estimated that 7% of sequences will belong to that species, and if species C was abundant, 29% of sequences would belong to that species (Figure 2). We note that conditioning the species-level random effect on abundance substantially improved model fit (Table 1), indicating that each species had a unique slope for different levels of rank-based abundance (Figure 3).

**Figure 2:**
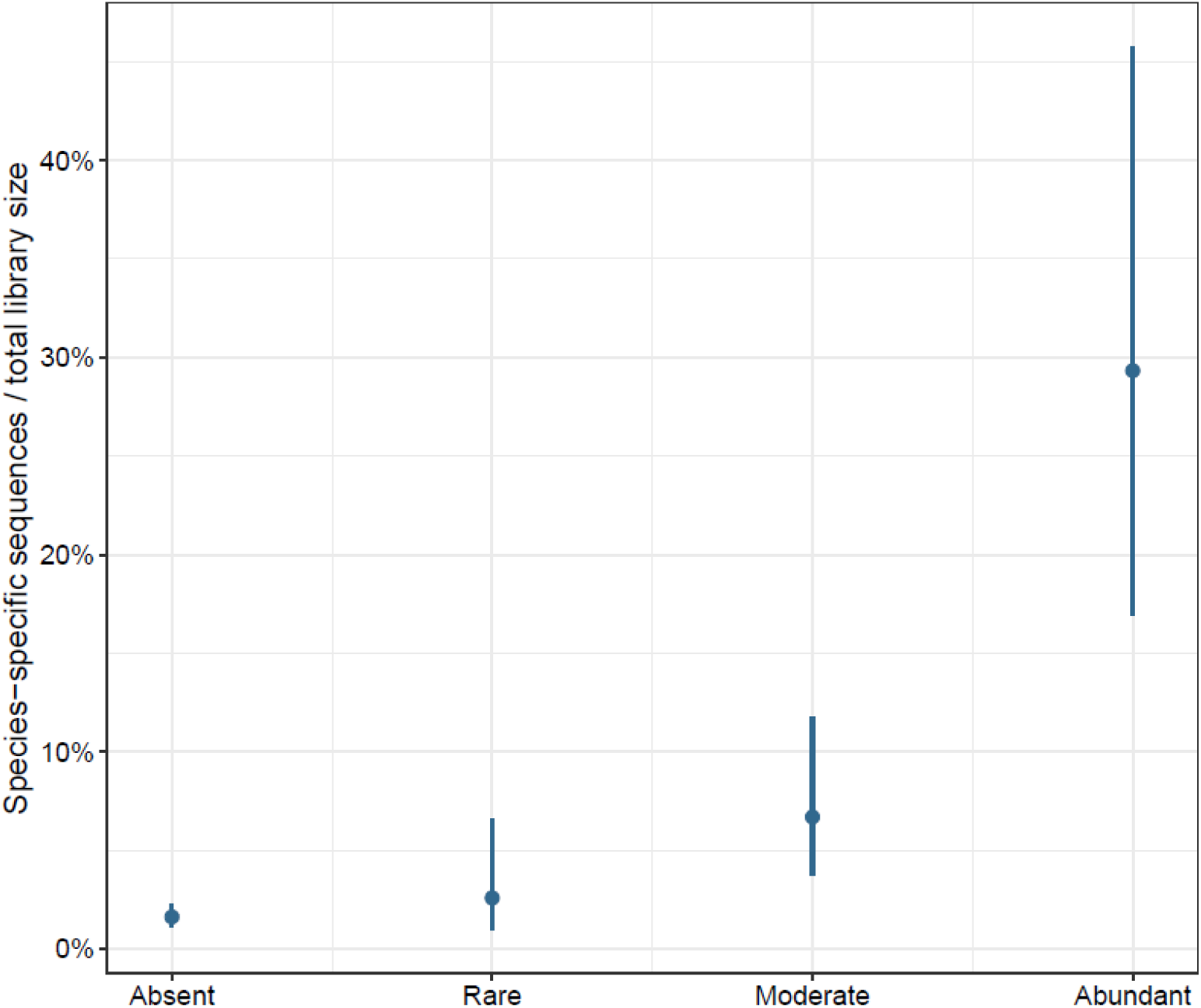
Back-transformed per-species read counts as a proportion of total library size by rank-based relative abundance category (absent, rare, moderate, abundant) across all species and all lakes in our study. For example, if a species is abundant in a lake, their species-specific read counts will be represented by 29% of the total library size of the samples in that lake. Species may occupy more than one relative abundance category (in different lakes). Error bars represent 95% confidence intervals.

**Figure 3:**
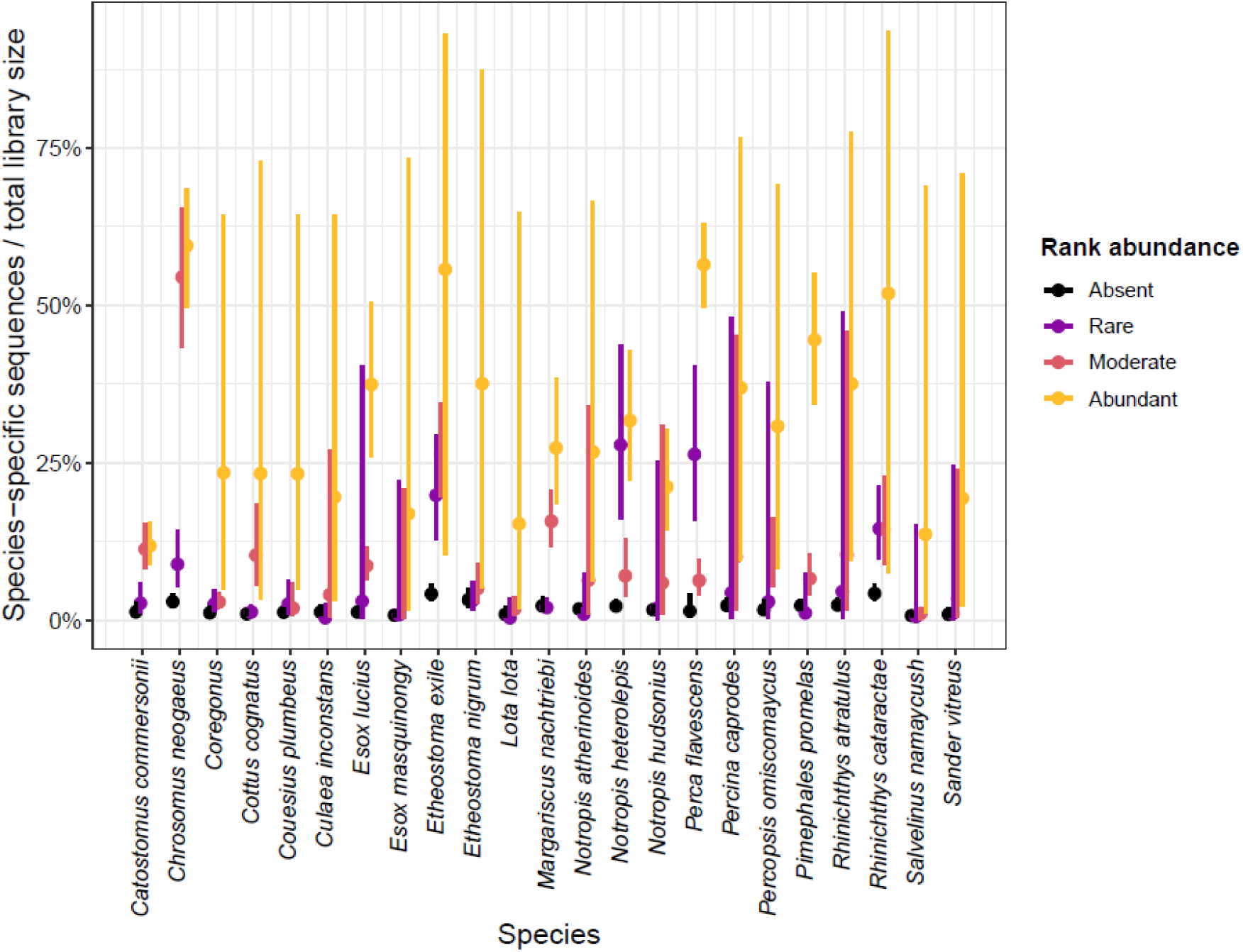
Marginal model predictions of rank-based abundance on species-specific sequence proportion of a sequencing library, conditional on the species-level random effect. The random slope model allowed the different levels of rank abundance to be different for each species. Error bars represent 95% confidence intervals; they are especially large for the levels of rank abundance which do not appear in the original dataset.

**Table 1:** Model AIC values for beta-binomial model comparing species-specific read count proportions to rank-based fisheries abundance estimates. The term listed in the first column is the term removed from the model to test against the ‘full’ model.

| Term tested (i.e., removed from model) | AIC | ΔAIC |
| --- | --- | --- |
| Full Model | 29004.83 | - |
| Abundance Dispersion term | 29016.72 | 11.89 |
| Abundance Zero-inflation term | 29018.46 | 13.63 |
| Random slope of species with rank<br>abundance | 29206.03 | 201.2 |
| Abundance fixed-effect term | 29024.91 | 20.08 |

Mean body mass data were available for 19 species across 20 lakes. Body mass for species represented in the dataset ranged from 0.2 g (*Etheostoma exile*) to 1859.8 g (*Esox lucius*), with a mean across populations and species of 304.4 g. The full model including an overdispersion term conditioned on abundance and an abundance term included in the zero-inflated component of the model failed to converge successfully. When either term was included individually, model convergence was successful, with the model including an overdispersion term conditioned on rank abundance exhibiting the lowest AIC (Table 2). We found a significant negative effect of mean body mass on the proportion of eDNA reads recovered (Figure 4). The proportion of reads recovered significantly decreased with increased mean body mass of the population (β = -0.0007, logit-scale; *p* < 0.001) indicating that, after controlling for relative abundance, larger-bodied species tended to produce less eDNA compared with smaller ones.

**Figure 4:**
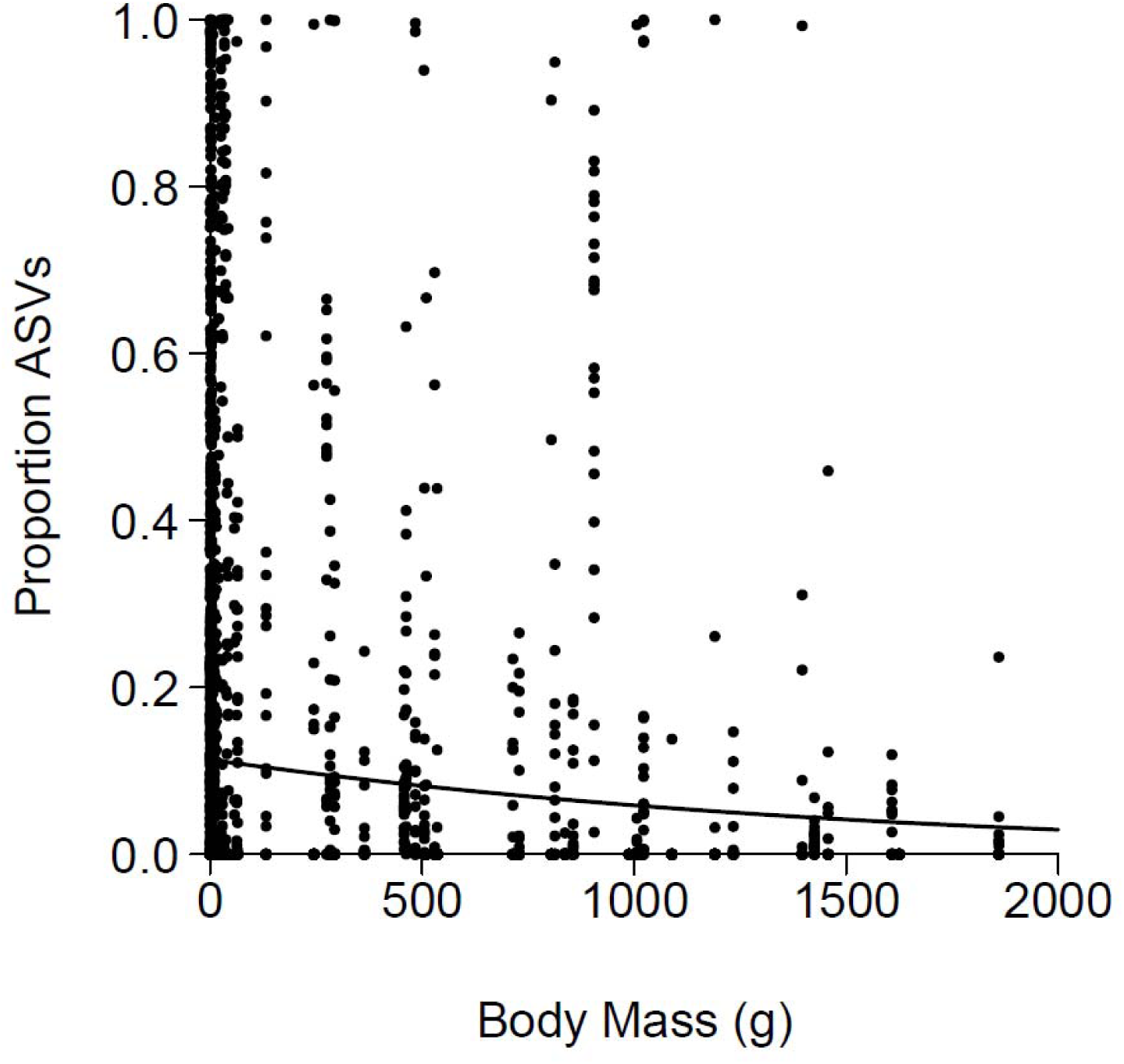
After abundance had been accounted for, there was a significant negative relationship between body mass of fish and the proportion of species-specific ASVs in the sequencing libraries, thought to occur due to an effect of allometry.

**Table 2:** Model AIC values for beta-binomial model comparing species-specific read count proportions to rank-based fisheries abundance estimates for the subset of species with mean body mass data available. The Abundance Dispersion term and Abundance Zero-inflation term were tested simultaneously due to the failure of the full model to converge. The following lower-order terms listed in the first column refer to the term removed from the model to test against the best-fit of the previously described two models.

| Term tested (i.e., removed from model) | AIC | ΔAIC |
| --- | --- | --- |
| Full Model* | NA | NA |
| Abundance Dispersion term** | 20564.2 | 2.9 |
| Abundance Zero-inflation term** | 20561.3 | - |
| Random slope of species with rank abundance | 20731.0 | 169.7 |
| Abundance fixed-effect term | 20577.2 | 15.9 |
| Body mass fixed-effect term | 20574.1 | 12.8 |

\*Model failed to converge
\*\* Models tested simultaneously

## Discussion

We found a highly significant relationship between within-species rank abundance and metabarcoding sequences from eDNA sampling taken across different habitats. This finding is based on robust replication, examining 18 taxa collected across 20 Boreal lakes in northwestern Ontario, using an average of 20 eDNA samples per lake. This finding confirms a long-held assumption that metabarcoding can be used semi-quantitatively to distinguish between sites in which the species is rare and those where it is abundant. While many studies have reported a positive relationship between fish abundance or biomass and eDNA sequences, 81% of 63 studies surveyed by Rourke et al. (2022) focused on single-species detection, with only eight field studies using eDNA metabarcoding to detect multiple species. Our study offers a key benefit by enabling rank-based abundance assessment of multiple species from a single metabarcoding dataset using the application of mixed effect models. In addition to traditional fisheries management approaches that focus on single species such as trout or walleye, this is a unique opportunity to assess multispecies habitats (including smaller fish species which are traditionally missed by gillnetting) in a semiquantitative manner with relatively low amounts of additional effort.

Crucially, our model supports the analysis of multiple within-species relative abundance across samples, rather than assessing the relative amounts of different species within a single lake. This type of methodology would be suitable for initial site assessments within an environmental impact assessment framework such as Phase 1 Habitat Assessment in the UK. Semi-quantitative schemes like the DAFOR scale (D = dominant, A = abundant, F = frequent, O = occasional, R = rare; Morris & Therivel, 2001) are commonly used by experienced biologists to rapidly assess species numbers or relative proportion during comparative, time-limited surveys such as vegetation surveys (Strong & Johnson, 2020).

Although these broad categories lack precision, the breadth allows a high degree of repeatability between users. Here, the rank abundance metric used has been validated on independently-collected abundance data from lakes with long-term sampling data. These abundance metrics have been applied successfully to fish species across the broad range of lakes in the IISD ELA region over the past 50+ years and have frequently been used to select new experimental or reference systems for whole lake experiments. Other implementations of abundance measurement rely on categories of logged numbers of individuals; e.g., the Whalley Hawkes Paisley Trigg method for recording river invertebrates under the Water Framework Directive in groups of 1-9, 10-99, 100-999, and >1000. As these categories are broad, it may be that with further investigation they could also be calibrated for across-site analysis with eDNA.

The relationship between eDNA sequence counts and abundance has previously been difficult to disentangle using metabarcoding methodologies. The success of our approach could result from analysis of relatively similar communities using consistent methodology over a moderately large number of habitats (n = 20 lakes), similar to Li et al., (2021) who studied anuran diversity across 71 waterbodies. Power analysis in a previous study that focused on winter shoreline sampling suggests that 10 samples are sufficient to capture 95% of fish species richness in 90% of 101 lakes (Sellers et al., 2024). Collection of a large number of samples may have helped to account for potential heterogeneity of detection in different parts of lakes, such as littoral habitats and the hypolimnion, as the lakes were highly stratified (Hänfling et al., 2016; Littlefair et al., 2023a; Yates et al., 2021). Additionally, the inclusion of lake as a random effect in our analysis accounted for the fact that certain lakes might have differing hydro-morphological or chemical conditions which may influence the amount of sequences recovered from the water (Caza-Allard et al., 2021; Klymus et al., 2015). This methodology may explain the difference between the strong relationship which we find, compared with the weak relationship between input biomass/DNA and sequence counts previously found in comparisons across species considering only single samples or habitats (e.g., Lamb et al., 2019). Despite this, there are implementation issues that require further development; for example, eDNA signal was present in lakes where the fish species was categorized as “absent”. This could be due to undersampling by fisheries-based assessment of small or rare species which can be missed by gillnets. Alternatively, the connected nature of the freshwater habitats can lead to downstream transport of small amounts of eDNA at the IISD ELA (Littlefair et al., 2023a), suggesting that the presence of signal in a lake where a species might be deemed absent could also be due to between-lake transport (Barnes & Turner, 2016; Deiner et al., 2016; Rourke et al., 2022).

By modelling species as a random slope, we allowed levels of sequence counts to vary across rank abundance categories for different species, which may have helped to account for any potential primer bias or species-specific eDNA shedding rates. There was substantial variation in the model predictions of the proportion of sequences belonging to different species, particularly for the “abundant” levels, which is why modelling species conditioned on abundance as a random slope rather than as a random intercept was a much better fit to our data. This variation in abundance levels could reflect either genetic mismatches between primer and target sequences or species-specific differences in ecological activity (Deiner et al., 2017). Differential eDNA shedding rates are thought to originate from variation in metabolic or seasonal activity such as spawning (Sassoubre et al., 2016; Troth et al., 2021).

We sampled during the summertime (July) and it is interesting to note that autumn and winter spawning fish such as the coregonids, burbot (*Lota lota*), and lake trout (*Salvelinus namaycush*) had the lowest predictions for the proportion of their sequences, whereas spring spawners such as Iowa darter (*Etheostoma exile*), yellow perch (*Perca flavescens*), longnose dace (*Rhinichthys cataractae*), and finescale dace (*Chrosomus neogaeus*) had some of the highest (Table S1). Other studies have found a similar effect of increased copy numbers during spawning (Bylemans et al., 2017; Thalinger et al., 2019; Tillotson et al., 2018), although few studies have examined a mixed-species approach with metabarcoding.

After accounting for abundance levels, we found a small but consistent negative effect of species body size on sequences recovered across our study lakes. Environmental DNA production is likely intrinsically related to key physiological rates (e.g., egestion and excretion) (Brown et al., 2004; Yates et al., 2021), and as organisms increase in mass their mass-specific metabolic and physiological rates tend to decline (Brown et al. 2004). These processes operate at both intra- (Jerde et al., 2019) and inter-specific levels (Vanni & McIntyre, 2016) and previous studies have found potential allometric effects on the recovery of eDNA at both an intra- (Maruyama et al., 2014; Yates et al., 2021) and inter-specific level (Stoeckle et al., 2021; Yates et al., 2023). While ontogenetic shifts in metabolic rates within and across species could account for these trends, it is also possible that other ecological factors could affect this relationship. Large-bodied fish, for example, tend to be piscivorous and carnivores tend to consume (and thus excrete) less per unit of body mass relative to herbivores and detritivores. A positive relationship between the degree of piscivory and body size across species could therefore also partially account for this pattern.

Various molecular and statistical methods are emerging for estimating abundance from eDNA data which account for the various sources of pipeline noise, such as primer bias towards certain species (Luo et al., 2023). Techniques like qPCR and ddPCR have shown promising results for single-species, across-site designs, with ddPCR enabling absolute quantification of DNA copy numbers (Hindson et al., 2011). The majority of published studies on fish eDNA have used these single-species methodologies, of which most have reported a positive relationship to abundance or biomass (Rourke et al., 2022). There are far fewer studies that consider the abundance of multiple species using metabarcoding, especially in a field context, although see Li et al. (2019); Pont et al. (2018); and Sard et al. (2019) for successful examples. Emerging approaches for estimating within-species abundance from metabarcoding data include the use of internal standards (“spike-ins”) during high-throughput sequencing. This involves the addition of the known abundance of a unique molecule that is added to all the samples. Absolute abundance within-species across-samples can be estimated by dividing the library size of the sample by the number of reads in each sample’s internal standard (Harrison et al., 2021; Luo et al., 2023). Similarly, the addition of unique molecular identifiers has been developed for absolute abundance estimation, in which sequences are randomly labelled with unique tags during library preparation (Hoshino & Inagaki, 2017).

The absolute number of copies of the gene in the original sample can be estimated by counting the number of unique molecular identifiers acquired by each species. Others have proposed to focus on eliminating the bias that obscures abundance information at particular stages of the molecular pipeline, for example by using mock communities to estimate different amplification rates (Shelton et al., 2023). Estimating both within-site and within-species abundance is complex, although Bush et al., (2023) recently applied a multi-species hierarchical model to macroinvertebrate data to explore presence-absence as a predictor of abundance to achieve this. However, this study used homogenate from bulk samples rather than eDNA, thus eliminating some issues resulting from variation in eDNA shedding.

The capacity of eDNA-based data to estimate abundance will always be called for with enthusiasm because abundance is a critical component of assessing the health, productivity and function of ecological communities. While many methodologies have been proposed, some require newly generated data (such as adding internal standards to sequencing libraries) or are impractical for large-scale, multi-species surveys due to cost or laboratory effort. Here we provide a pragmatic but efficient analysis to determine the rank-based abundance of multiple species, suitable for practical application in a fisheries management context.

## Acknowledgements

Many IISD ELA researchers and students helped to generate the field data for which we are grateful. Sonya Michaleski and Rachel Henderson contributed specific field assistance to this project. Collection of historical data contributing to fish abundance metrics was led by Ken Mills, Sandra Chalanchuk and Doug Allan. We thank Richard Nichols and Maxwell Farrell for statistical advice. This work was funded by a Mitacs Accelerate award (JEL, MEC), NSERC Discovery grants (MEC, MDR), Québec Centre for Biodiversity Science Excellence award (JEL), the IISD ELA, the WSP Montréal, Environment Department, a UKRI Future Leaders Fellowship (JEL), and Canada Research Chair awards (MDR and MEC).

## Author contributions

JEL, MDR, and MEC conceptualised the study and contributed to experimental design. LDH and MDR contributed datasets and analysis of the fish catch and abundance data. JEL, MDR and LDH performed field work, JEL performed molecular lab work and JEL and MDR performed statistical analysis and visualisations. JEL wrote the first draft of the manuscript and all authors contributed to editing and writing subsequent drafts.

## Data availability statement

Data (Littlefair et al., 2023b) are available from Dryad: https://doi.org/10.5061/dryad.zs7h44jd0. Code and three dataframes specific to this project are available during peer review, which will be made public on manuscript acceptance: https://figshare.com/s/58072c2d3eab18302245

