## Supporting Information for "A strong relationship between environmental DNA metabarcoding and rank-based abundance of fish"

| Page | Item |
| --- | --- |
| 2 | Summary of molecular survey methods applied to eDNA from the IISD ELA lakes |
| 5 | Validation of rank-based abundance with standardized (measured and estimated) abundance data |
| 8 | Figure S1: Relationship between rank abundance and CPUE or mark-recapture data |
| 10 | Table S1: Fish species used in this study, alongside spawning season and method of generating body mass data |
| 13 | References used in the Supporting Information |

### Summary of molecular survey methods applied to eDNA from the IISD ELA lakes

### Methods

*eDNA sampling*

The sampling strategy described in full in Littlefair et al., (2023) was applied to each lake, which is briefly summarised here for the coherence of this manuscript. Pelagic surface samples (500ml each) were taken along a transect at five evenly spaced intervals across each lake, including the deepest point of the lake (n = 10). In addition, deep water samples (2m from the sediment surface) were taken at sampling stations #3 and #4 on the transect using a van Dorn bottle, (n = 2). We also took two samples from the shoreline of each lake, and two samples from major inflows and outflows that could be identified on each lake. In total, 386 samples were used in this study. Water samples were filtered within six hours of collection using 47mm GF/F filters (ThermoFisher Scientific; nominal pore size = 0.7µm). Filters were dry frozen at -20⁰C and transported to McGill University on dry ice for molecular analysis.

*eDNA molecular analysis*

DNA was extracted from filters using the Qiagen Blood and Tissue kit with some modifications to the manufacturer’s instructions. 370µl buffer ATL was used in the initial incubation step, filters were incubated in ATL and proteinase K for 16 hours overnight, and the DNA was eluted in 2 x 40µl of AE buffer and stored at -80⁰C after elution. Extractions were treated with the OneStep PCR Inhibitor Removal Kit (Zymo Research, Irvine, California).

We targeted fish biodiversity using the MiFish-U (Miya et al., 2015) primers which amplify part of the 12S region. DNA was amplified in triplicate reactions using 7.4µl nuclease free water (Qiagen), 1.25µl 10X buffer (Genscript), 1mM MgCl_2_ (ThermoFisher Scientific), 0.2mM GeneDirex dNTPs, 0.05mg bovine serum albumen (ThermoFisher Scientific), 0.25mM each primer, 1U taq (Genscript) and 2µl DNA in a final volume of 12.5µl. The thermocycling protocol involved denaturation at 94°C for two minutes, followed by 50 cycles of five seconds at 98°C, 10 seconds at 50 °C and 10 seconds at 72°C, followed by a final elongation at 72°C for five minutes. Triplicate PCR amplicons for each sample were combined, cleaned with AMPure beads and indexed with the Nextera DNA indexing kit for 96 samples (Illumina). A second clean-up with AMPure beads was performed, and libraries were quantified and normalised to 5ng/μl. Equimolar amounts of DNA were combined and samples were allocated across five sequencing lanes and sequenced with even depth per sample. Sequencing was conducted using 2x300bp Illumina MiSeq at the McGill University and Génome Québec Innovation Centre, Montréal.

To prevent contamination, we processed the samples in a clean, pre-PCR dedicated laboratory. Before beginning any work, we thoroughly cleaned benches with 10% bleach. Laboratory equipment was cleaned with 70% ethanol and RNAse wiper. Negative controls were included at each major step: field sampling, DNA extraction, and PCR amplification. A combined negative control was sequenced, even in the absence of visible gel bands.

*Bioinformatics*

We used a denoising pipeline to filter errors and cluster sequences into amplicon sequence variants or ASVs (Callahan, McMurdie, & Holmes, 2017). Samples were received as demultiplexed fastq files from Génome Québec. Nonbiological nucleotides (primers, indices and adapters) were removed using cutadapt (Martin, 2011). Paired reads were merged using PEAR with a minimum quality score threshold of 25 and a minimum length of 100 bp after trimming (Zhang, Kobert, Flouri, & Stamatakis, 2014). Quality scores for sequences were analysed with FASTQC (Andrews, 2010). Reads were length filtered between 152–192 bp (± 20bp around the target amplicon size). Amplicon sequencing variants were generated using the UNOISE3 package (Edgar, 2016), which uses a denoising pipeline to remove sequencing error and to cluster sequences into single variants (100% similarity). The full bioinformatic pipeline is available from <https://github.com/CristescuLab/YAAP>.

We assigned taxonomy to the ASVs using BLAST+ (Camacho et al., 2009) with high stringency parameters (98% identity, 90% query coverage) and used the last common ancestor algorithm in BASTA (Kahlke & Ralph, 2019) to assign taxonomic identity. We matched to a local database composed of sequences from fish species during the last 50 years of monitoring data at the IISD-ELA as well as government surveys of the same region. We performed an adjustment of sequence numbers based on the small amount of contamination that appeared in the mock community; this was done by calculating proportions for each species that would remove the contamination from the mock community and the negative controls and then adjusted this to every sample’s library size. We then subtracted this number of sequences from every library. We then grouped together ASVs which matched to the same fish species, as populations might have more than one haplotype (no ASV matched to more than one fish). *Chrosomus eos* and *Chrosomus neogaeus* form mitochondrial hybrids in this area so this ASV could only be assigned to *Chrosomus* genus (Mee & Taylor, 2012). *Coregonus artedi* and *Coregonus clupeaformis* are closely related and could only be identified to genus level (Littlefair et al., 2023). For the rank-based abundance assignment we retained the highest rank abundance for the genus where the two species appeared in the same lake.

### Validation of rank-based abundance with standardized (measured and estimated) abundance data

### Methods

A general metric of relative abundance (besides mark-recapture and catch-per-unit-effort estimates) that has been implemented at the IISD-ELA since its inception has been the semi-quantitative rank abundance method of Beamish et al. (1976). This assigns qualitative descriptors of 0 (absent), R (rare), M (moderately abundant) and A (abundant) to species across lakes. This was initially performed based primarily on initial surveys conducted during 1972-73 at the Experimental Lakes Area using gillnets, trapnetting and minnow trapping (for details see Beamish et al. 1976). Similar methods have been applied to other lakes outside of the original survey by Beamish et al (1976) to establish similar metrics across additional lakes. However, the precise means by which these estimates were made or how categories were set is not documented, and therefore requires validation using independently collected quantitative measures of abundance collected from these lakes since these original surveys.

To evaluate the usefulness of the Beamish et al. (1976) ranked designations as a quantitative measure of fish abundance and, therefore, whether it is an accurate marker of abundance for comparison with quantitative eDNA estimates, we compared the original rank abundance assignments (0 = absent, R = rare, M = moderately abundant, A = abundant; Beamish et al., 1976) to several lakes, which have been subjected to annual quantitative surveys.

We examined data from lakes 114, 189, 191, 222, 223, 224, 226, 239, 305, 378 and 382, as listed in the original Beamish et al. (1976) report, as they had one or more overlapping fish species in multiple rank abundance categories. To cover each of the abundance categories for each fish species investigated, we also included data from lakes 373 and 260; these lakes were first surveyed in the early 1980s by at least one staff member who was included in the original Beamish survey (Ken Mills, Research Scientist Emeritus, Department of Fisheries and Oceans Canada), ensuring that categorical classifications of abundance were performed consistently with the original surveys. For lakes that experienced ecosystem manipulations affecting fish densities (i.e., field experiments in lakes 191, 222, 223, 226, 305), time periods over which manipulations occurred were excluded from the analysis.

The following species from the original Beamish et al. (1976) report were used to assess the accuracy of the original assignments; Lake Trout (*Salvelinus namaycush*), White Sucker (*Catostomus commersonii*), Fathead Minnow (*Pimephales promelas*), Northern Pearl Dace (*Magariscus nachtriebi*), Northern Redbelly Dace (*Chrosomus eos*), and Slimy Sculpin (*Cottus cognatus*). All of these species are also detected by eDNA in the present study.

Quantitative assessments of fish abundance at the ELA are performed primarily in spring and fall. Therefore, we chose the season of data to evaluate relationships with rank abundance to correspond with the predominant breeding period for the particular species of interest (Table S1; i.e., spring for White Sucker, Northern Redbelly Dace, Northern Pearl Dace, Slimy Sculpin; autumn for Lake Trout, Fathead Minnow). Though fathead minnow spawn several times during the late summer, the fall (Aug-October) sampling period was chosen to assess this species to capture cumulative spawning efforts over the previous open water period. Fish were collected historically under the authority of Fisheries and Oceans Canada, and contemporary collections were made under the permission of the Canadian Council for Animal Care by Lakehead University (Animal User Protocol #1464656) and the Ontario Ministry of Natural Resources and Forestry (License to Collect Fish for Scientific Purposes #1085769).

With the exception of Lake Trout, all species listed above were assessed for quantitative relative abundance using catch per unit effort (CPUE) from Beamish-style trap nets (Beamish 1973). Typically, multiple trap nets are fished in a lake for several days at a time during a given season of sampling. The catch (number) for each species is standardized by the number of nets and number of days for each visit to the lake (i.e. number per net*days). This value was averaged over all sampling events in a given sampling period (e.g. spring or autumn of each year the lake was sampled). Lake trout individuals are tagged upon capture in the spring and fall, and abundance for this species is assessed using a mark-recapture model that allows for births and deaths in the population (using the POPAN module in program Mark, implemented through RMark; R version 4.1.2, R Core Team, 2021). Using these methods, means of annual population estimates for each lake (outside of manipulated periods where relevant) were estimated. In order to analyze these different abundance metrics on a similar scale across all species, all abundance metrics for each species were Z-score standardized (centred using the season-specific mean abundance measure where appropriate, and divided by the appropriate standard deviation).

We used a linear mixed effect model framework (using the *lme4()* package in R) to evaluate whether there was a general pattern in measured abundance with rank abundance estimates across all species. We included species as a random intercept in our models. Because a pattern of heteroscedasticity (increasing variance with rank abundance) was observed in our original linear mixed effects model, we proceeded with a generalized linear mixed effects model with a Gamma distribution and a log link function, which removed any patterns in residual plots from the model fit. To facilitate model fit under a Gamma distribution, standardized abundance values were rendered positive by adding 1.1 to all values. We used the *lmerTest()* package to determine significance of main effects (using the Satterthwaite approximation of degrees of freedom), and the *glht()* function of the *multcomp()* package (with *P*-values adjusted using the ‘holm’ method) to assess differences in measured abundance between rank abundance categories. Conditional and marginal *R*^2^ values for the fixed effect of rank abundance were estimated using the r2_nakagawa function in the *performance()* package. Bootstrapped 95% confidence intervals for predicted values of each level of rank abundance were estimated for data plots using the *bootMer()* function with 999 bootstraps.

### Results

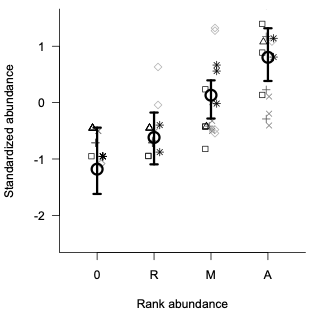
The semi-quantitative rank abundance classification significantly explained the variation we observed in relative or absolute abundance across all fish species (*F_3,65_* = 33.0, *P*<0.0001; conditional *R*^2^ = 0.62; marginal *R*^2^ = 0.59). A post-hoc test demonstrated that all classes were significantly different from one another, in the order expected (0<R<M<A; Figure S1).

Figure S1: Predicted means (large circles) and bootstrapped 95% confidence intervals (error bars) from linear mixed effect model demonstrating correspondence between rank abundance assigned by Beamish et al. (1976) with a standardized measure of fish abundance (catch per unit effort or mark-recapture abundance estimates, see text). Squares are White Sucker, upward triangles are Northern Redbelly Dace, plus signs are Fathead Minnow, crosses are Northern Pearl Dace, diamonds are Slimy Sculpin, stars are Lake Trout.

Table S1: Spawning seasons for the fish species found at the IISD ELA, North West Ontario. Spring refers to March-May, Summer refers to June-August, Autumn refers to September-November and Winter refers to December-February. Body mass indicates how body mass was derived for each species for the analysis considering mean weight of species in each lake (see Methods for details).

| **Latin name** | **Common name** | **Spawning season** | **Body mass** |
| --- | --- | --- | --- |
| *Catostomus commersonii* | White Sucker | Spring | Observed |
| *Chrosomus eos* | Northern Redbelly Dace | Summer | Estimated from length^2^ |
| *Chrosomus neogaeus* | Finescale dace | Spring | Estimated from length^2^ |
| *Coregonus artedi* | Lake Herring | Autumn + Winter | Observed |
| *Coregonus clupeaformis* | Lake Whitefish | Autumn | Observed |
| *Cottus cognatus* | Slimy Sculpin | Spring | Observed or estimated from length^3^ |
| *Couesius plumbeus* | Lake Chub | Summer | Estimated from length^4^ |
| *Culaea inconstans* | Brook stickleback | Spring | Estimated from length^5^ |
| *Etheostoma exile* | Iowa darter | Spring | Estimated from length^6^ |
| *Etheostoma nigrum* | Johnny darter | Spring | Estimated from length^2^ |
| *Esox lucius* | Northern Pike | Spring | Observed |
| *Esox masquinongy* | Muskellunge | Spring | Not used |
| *Lota lota* | Burbot | Winter | Observed |
| *Margariscus nachtriebi* | Northern Pearl dace | Spring | Estimated from length^2^ |
| *Notropis atherinoides* | Emerald Shiner | Summer | Estimated from length^2^ |
| *Notropis heterolepis* | Blacknose Shiner | Spring + Summer | Estimated from length^7^ |
| *Notropis hudsonius* | Spottail Shiner | Summer | Estimated from length^2^ |
| *Perca flavescens* | Yellow Perch | Spring | Estimated from length^1^ |
| *Percina caprodes* | Logperch | Spring | Not used |
| *Percopsis omiscomaycus* | Trout-perch | Summer | Mean value from other ELA lakes^8^ |
| *Pimephales promelas* | Fathead Minnow | Summer | Estimated from length^9^ |
| *Rhinichthys atratulus* | Blacknose dace | Spring | Not used |
| *Rhinichthys cataractae* | Longnose Dace | Spring + Summer | Estimated from length^4^ |
| *Salvelinus namaycush* | Lake Trout | Autumn | Observed |
| *Sander vitreus* | Walleye | Spring | Not used |

Equations used for mass estimations for length: ^1^Hayhurst et al. 2018 (Lake 239, 2016 relationship); ^2^Mushet et al. in review; ^2^Schneider et al. 2000; ^4^previously unpublished data, equations provided below; ^5^Estimated based on equation across two source references (Devine et al. 2005; Yamamoto et al. 2022), with default weighting provided for nine-spine stickleback collected from Lake Superior, as reported in fishbase.org (Froese and Pauly 2024); ^6^Based on equation provided for blackside darter reported in Schneider et al. 2000; ^7^Based on equation provided for spottail shiner reported in Schneider et al. 2000; ^8^Based on mean of observed mass values from Lake 421 at the IISD ELA; ^9^Milling et al. 2020.

Previously unpublished length-weight relationships for IISD ELA fishes

In order to estimate mass from length, weight-length relationships were developed using previously unpublished IISD ELA data. Equations, along with associated statistical parameters are as reported below and derived from fork length measurements:

**Lake Chub (*Couesius plumbeus*):**

Log_10_(weight) = 2.2662*Log_10_(fork length) -4.2566, *F*_1,177_ = 868.3, *p* < 0.0001, *R*^2^ = 0.83

Derived from data across Lakes 223, 260 and 626 (with 13, 9 and 157 observations, respectively).

**Longnose dace (*Rhinichthys cataractae*):**

Log_10_(weight) = 3.3215*Log_10_(fork length) -5.5386, *F*_1,81_ = 500.4, *p* < 0.0001, *R*^2^ = 0.86

Derived from data across Lakes 305, 377 and 626 (with 2, 5 and 76 observations respectively).
